# Meningeal inflammation and arachnoid barrier breakdown in a mouse model of neonatal bacterial meningitis

**DOI:** 10.64898/2026.03.04.709573

**Authors:** Sol Kim, Luke R. Joyce, Amanda Brady, Brady L. Spencer, Bradley Pawlikowski, Julia Derk, Kelly S. Doran, Julie A. Siegenthaler

**Affiliations:** University of Colorado Anschutz, Department of Pediatrics, Section of Developmental Biology, Aurora, CO, USA; University of Colorado Anschutz, Cell Biology Stem Cells and Development Graduate Program, Aurora, CO, USA; University of Colorado Anschutz, Department of Immunology and Microbiology, Aurora, CO, USA; University of Virginia, Department of Microbiology, Immunology & Cancer Biology, Charlottesville, VA; Howard Hughes Medical Institute, The Collective for Psychiatric Neuroengineering, Durham, NC, USA; Duke University School of Medicine, Department of Psychiatry, Durham, NC, USA; North Carolina Central University, Department of Biological and Biomedical Sciences, Durham, NC, USA

**Keywords:** Group B *Streptococcus*, leptomeninges, Claudin-11, cytokines, border-associated macrophages, meningeal fibroblasts

## Abstract

Newborns are especially susceptible to bacterial meningitis, primarily caused by Group B *Streptococcus* (GBS), due to incomplete maturation of immune and barrier systems. While meningitis is well known to break down the blood-brain barrier (BBB), how the meningeal arachnoid barrier, a critical component of the blood-cerebrospinal fluid barrier (B-CSFB), responds to infection is poorly understood. Using a neonatal mouse model of bacterial meningitis, we demonstrate that GBS infection significantly increases arachnoid barrier permeability, coinciding with alterations in Claudin-11 tight junction distribution and elevated meningeal production of proinflammatory cytokines (IL-6, TNF-α, CXCL1). CD206^+^/Lyve1^+^ border-associated macrophages (BAMs) undergo significant morphological and molecular activation post-infection, but their depletion prior to GBS infection did not attenuate arachnoid barrier leakage or inflammatory cytokine levels during infection. We show that meningeal fibroblasts are a main source of proinflammatory cytokines in response to GBS infection and that exposure to the inflammatory cytokine TNF-α alone is sufficient to induce neonatal arachnoid barrier breakdown. These results support neonatal arachnoid barrier is vulnerable to cytokine-induced breakdown in bacterial infection and highlight the role of non-immune meningeal cells like fibroblasts during bacterial infection.

## Introduction

The central nervous system (CNS) is protected by essential barriers that regulate the entry of peripheral molecules, cells and pathogens. These barriers include the blood-brain barrier (BBB) at the level of CNS microvasculature and the blood-cerebrospinal fluid barrier (B-CSFB) at the choroid plexus and meninges (Engelhardt and Sorokin, 2009). CNS barriers can be compromised following CNS injury, infection or in neurological diseases, in which they exhibit structural and functional impairment that can exacerbate neuroinflammation and neurological dysfunction. One such example is bacterial meningitis, which occurs when blood-borne bacteria enter the leptomeninges, brain, and choroid plexus and induce uncontrolled inflammation (Huang et al., 2000; Kim et al., 2015; Schwerk et al., 2015; Wang et al., 2023; Xu et al., 2024). Neonatal bacterial meningitis—primarily caused by Group B *Streptococcus* (GBS) and *Escherichia coli* (*E. coli*)—is a leading cause of neurodevelopmental disorders worldwide. Occurring in roughly 0.3 per 1000 live births annually, these infections result in 5-20% mortality while 20-50% of survivors develop life-long neurological sequelae (Bundy et al., 2025; Edwards et al., 1985; Ku et al., 2015; Phares, 2008). As the primary etiology, GBS is a Gram-positive bacterium that first invades peripheral systems to induce bacteremia, with subsequent meningeal inflammation and BBB breakdown (Doran et al., 2005; Coureuil et al., 2017; Tavares et al., 2022). Newborn cases of bacterial meningitis tend to be particularly severe due to their immature immune system and barrier systems across the body (Deshayes De Cambronne et al., 2021; Huang et al., 2000; Kim et al., 2023a; Tavares et al., 2022; Travier et al., 2021a; Van Hove et al., 2019). Despite this, most studies modeling bacterial meningitis in animal models are adult- and BBB-centric. They often exclude developmental timepoints mirroring when humans are most susceptible and overlook the impact of meningitis on the B-CSFB, in particular the specialized meningeal barrier layer called the arachnoid barrier (AB) (Kim et al., 2023; Wang et al., 2023).

The AB is a key component of the B-CSFB and part of the meninges, a membranous structure surrounding the CNS that consist of three main layers: the dura, arachnoid and pia. Each layer of the meninges contains specialized fibroblast populations that mediate specific functions. The outermost layer below the calvarium is the dura, which contains fenestrated blood vessels, lymphatic vessels, and diverse immune populations. The inner two layers are collectively referred to as the leptomeninges; the brain-adjacent pia layer is composed of a single layer of fibroblasts and blood vessels with barrier properties, while the arachnoid layer contains the AB segregating the leaky dura from the barrier vasculature of the pia and the brain (Betsholtz et al., 2024; Derk et al., 2021). The AB is comprised of 2-3 layers of epithelial-like fibroblasts that are interconnected by a combination of tight (Claudin-11, ZO-1) and adherens (E-cadherin, VE-cadherin) junction proteins (Nabeshima et al., 1975; Pietilä et al., 2023; Mapunda et al., 2023). The AB is a key component of the B-CSFB, serving as a selective barrier that regulates the movement of molecules between the overlying dura and the underlying CSF-filled sub-arachnoid space. (Balin et al., 1986; Derk et al., 2023; Mapunda et al., 2023; Rodriguez-Peralta, 1957; Roth et al., 2013; Smyth et al., 2024; Zhang et al., 2022). Recent work in mice linked neuroinflammation with an induced AB leakage, altered AB junctional protein expression and impaired behavior (Zhang et al., 2022), supporting the importance of AB integrity in brain function. However, unlike the well-characterized breakdown of BBB junctional proteins during inflammation, how local inflammation impacts the AB integrity is just beginning to be examined.

CNS- or border-associated macrophages (BAMs) are a resident macrophage population at various CNS barriers (Goldmann et al., 2016; Vara-Pérez and Movahedi, 2025). BAMs are found in the meninges, choroid plexus, and perivascular spaces, and their activation has been implicated in various pathologies, including meningitis, ischemic stroke, Alzheimer’s disease, and multiple sclerosis (Kierdorf et al., 2019; Wang et al., 2023; Sun and Jiang, 2024; Van Hove et al., 2025). BAMs express the macrophage marker CD206 (encoded by *Mrc1)*, and BAMs in the leptomeninges (L-BAMs) are CD206^+^/Lyve1^+^ while dural BAMs are Lyve1 negative. Fibroblasts also play a crucial role in CNS inflammation and injury, and the pro-inflammatory response of meningeal fibroblasts has been shown to potentiate spinal cord injury, stroke, and multiple sclerosis phenotypes (Buechler et al., 2021; Dorrier et al., 2022; Cupovic et al., 2016; Duan et al., 2018; Bajénoff et al., 2006; Magliozzi et al., 2006; Pikor et al., 2015). Despite their prevalence and immunomodulatory capabilities, the roles of L-BAMs and meningeal fibroblasts during bacterial meningitis and influence on AB integrity have yet to be investigated.

In this study, we analyze the response of the AB and meninges in neonatal mice using a GBS meningitis model. We show that AB functional integrity is lost after exposure to GBS, which coincides with mislocalization of tight junction protein Claudin-11. We observed a strong proinflammatory response from both meningeal fibroblasts and L-BAMs and find that L-BAM depletion concurrent with GBS infection does not protect the AB from breaking down. Using the neonatal animal model and primary cultures, we detail leptomeningeal and dural fibroblast production of inflammatory cytokines post-infection which include IL-6, CXCL1 and TNF-α. Lastly, we show that inflammation induced by exogenous TNF-α alone is sufficient to breakdown the AB and alter Claudin-11 organization. These results demonstrate the vulnerability of neonatal AB to cytokine-induced breakdown in bacterial infection and highlight the significance of non-immune meningeal cells during bacterial infection.

## RESULTS

### AB functional integrity is altered in a murine model of neonatal bacterial meningitis

Although bacterial meningitis can occur in all ranges of age, newborns are especially susceptible to bacterial meningitis due to the immaturity of their physiological barriers and immune system (Huang et al., 2000; Kim et al., 2023b; Travier et al., 2021b). Previous studies have reported that early bacterial infection leads to compromised intestinal, choroid plexus, and endothelial barriers in the postnatal mouse brain (Huang et al., 2000; Travier et al., 2021b; Wang et al., 2023), yet the vulnerability of the AB has not been tested.

To quantify functional changes in neonatal AB during bacterial meningitis, we utilized a bacterial meningitis model in early postnatal mice using Group B *Streptococcus* (GBS) (Joyce et al., 2024) (see **Methods**; **Supp. Fig. 1A-C**). We challenged postnatal day (P)2 mice with either GBS (2×10^6^ CFU, strain COH1) or PBS (mock) and harvested them 18-24 hours post-infection (hpi) when most of the infected animals became moribund. To assess the functional integrity of the AB, we adapted a transcranial dye assay used to investigate AB integrity in adult mice (Zhang et al., 2022) (**Fig. 1A**; see **Methods**). In the assay, a fluorescent Biocytin-TMR dye (molecular weight: 870Da) is applied to the exposed calvarium. The dye passes through the porous bone and the dura, and in a control situation where the AB is intact, the tracer accumulates at the AB. However, if the integrity of the AB has been compromised, the tracer passes through the subarachnoid space and into the superficial cortex. The resulting fluorescent tracer signal in the superficial cortex can be quantified using microscopy analysis of thick sections or with a plate reader-based fluorescence quantification using dissected cortical tissue. We analyzed Biocytin-TMR signal in the superficial cortex using both methods and found that the tracer had significantly diffused into the superficial cortex of GBS-infected mice by 18-24 hpi, while the dye was confined in the leptomeninges (LPM) of mock-infected animals (**Fig. 1B, C**). This data shows that the AB integrity is lost following bacterial infection in neonatal mice.

**Figure 1.**
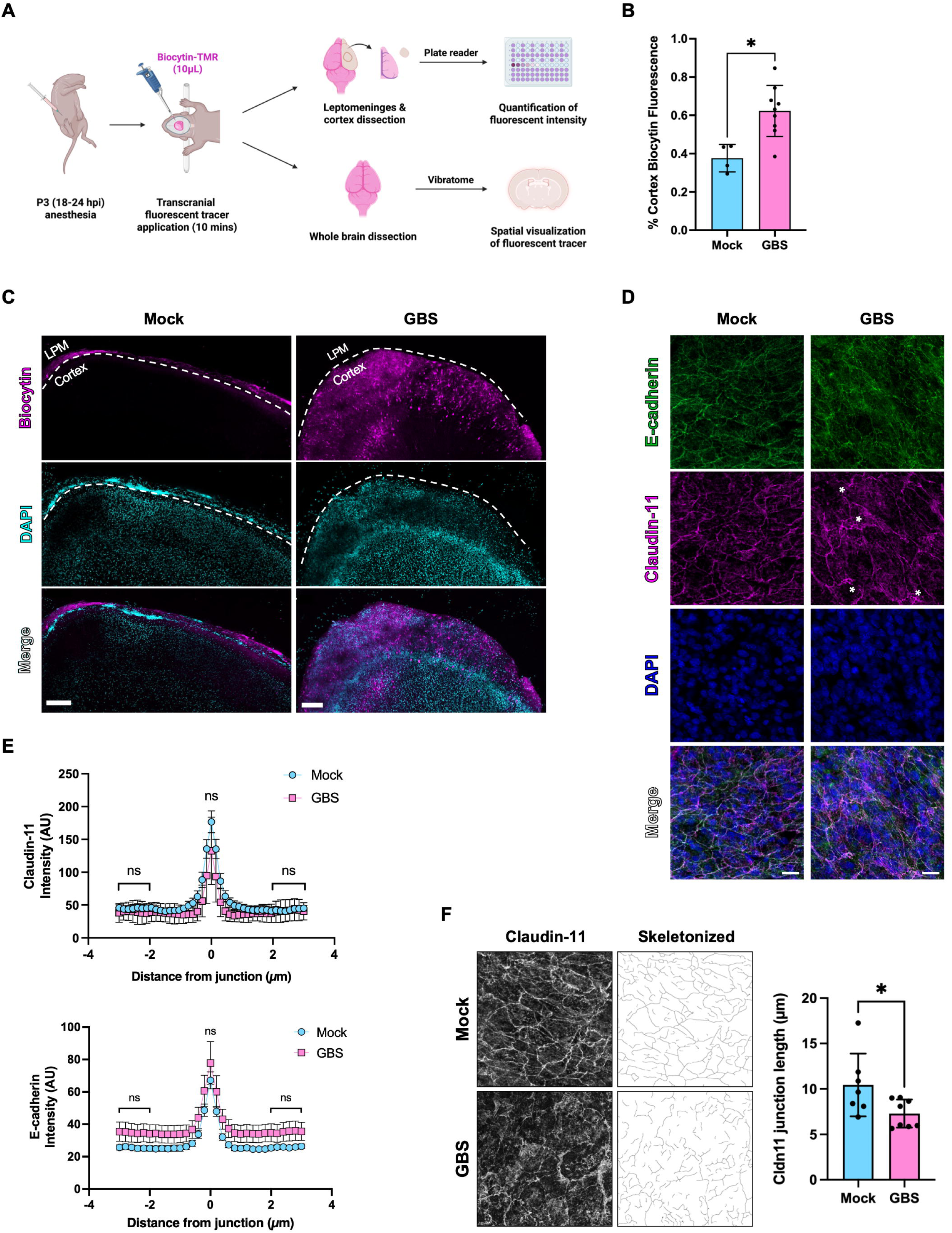
Neonatal arachnoid barrier is disrupted during GBS infection. **(A)** Schematic of arachnoid barrier integrity assay (ABIA) using a plate reader and thick section imaging (adapted from Zhang et al., 2022). **(B)** Plate reader quantification of % Biocytin-TMR fluorescence in the superficial cortex (cortical fluorescence normalized to LPM fluorescence). (n=4 mock, n=9 GBS) **(C)** Visualization of Biocytin-TMR in the leptomeninges (LPM) and cortex between mock-infected and GBS-infected group (500µm coronal brain sections). **(D)** Representative images of E-cadherin^+^ (E-cad) and Claudin-11^+^ (Cldn11) AB junctions on LPM whole mounts using immunofluorescence (IF). Asterisks indicate locations of Cldn11^+^ junction aberrance. **(E)** Line intensity analysis of E-cad^+^ and Cldn11^+^ junctions (peak at +0-0.45µm) and non-junctions (between +2-3µm) on LPM whole mounts. **(F)** Ridge Detection analysis of Cldn11+ junctions (left) and quantification (right) on LPM whole mounts. (n=7 mock, n=8 GBS for 1E and 1F) Statistics: Mann-Whitney *U* test, * = p < 0.05, ns = not significant (p-value > 0.05); mean and SD. Scale bars = (C) 200µm, (D) 20µm.

The AB cells are held together by extensive junctional proteins, two of which are the adherens junction E-cadherin (E-cad) and the tight junction Claudin-11 (Cldn11) (Pietilä et al., 2023; Smyth et al., 2024). We next conducted an immunofluorescence (IF) analysis of AB junctional proteins using LPM whole mounts, where the LPM are dissected off the surface of the brain and processed for staining (Jones et al., 2022). Quantitative analysis of E-cad and Cldn11 expression at AB cell junctions using a fluorescence intensity profile showed that both E-cad and Cldn11 were still present at cell junctions following infection and that E-cad^+^ junctions appeared normal (**Fig. 1D, E; Supp. Fig. 1D**). In contrast, Cldn11^+^ junctions had a discontinuous appearance, with a significantly lower average length and more cytoplasmic signal in GBS-infected group compared to mock (**Fig. 1F; Supp. Fig. 1E**). Taken together, these data demonstrate that the AB becomes leaky and Cldn11^+^ tight junctions are altered after infection.

### GBS infection induces local proinflammatory response in the meninges

Disruption of junctional proteins is a common indicator of barrier breakdown in physiological systems upon inflammation, including the BBB and choroid plexus epithelium (Kim et al., 2015; Castro Dias et al., 2019; Cui et al., 2020; Solár et al., 2020; Galea, 2021; Xu et al., 2024). A major driver behind junctional disruption is proinflammatory cytokines and chemokines such as TNF-α, IL-6, IL-1β and IFN-γ (Gryka-Marton et al., 2025; Walsh, 2000; Yang et al., 2019). To assess the inflammatory response in our GBS infection model, we measured the levels of common proinflammatory molecules in dura, LPM, and brain using a multiplex cytokine assay. The results showed a significant increase in IL-6, CXCL1, TNF-α, IL-1β, and IL-10, post-infection in dura, LPM, and brain (**Fig. 2A**). Consistent with systemic nature of the infection model, there was increased expression of these cytokines in peripheral organ (liver) and plasma (**Supp. Fig. 2A**). Dura showed the highest post-infection levels but the levels in LPM and brain were also significantly higher post-infection (**Fig. 2A, B**). These data show that GBS infection triggers a robust inflammatory signaling response in the meninges and brain.

**Figure 2.**
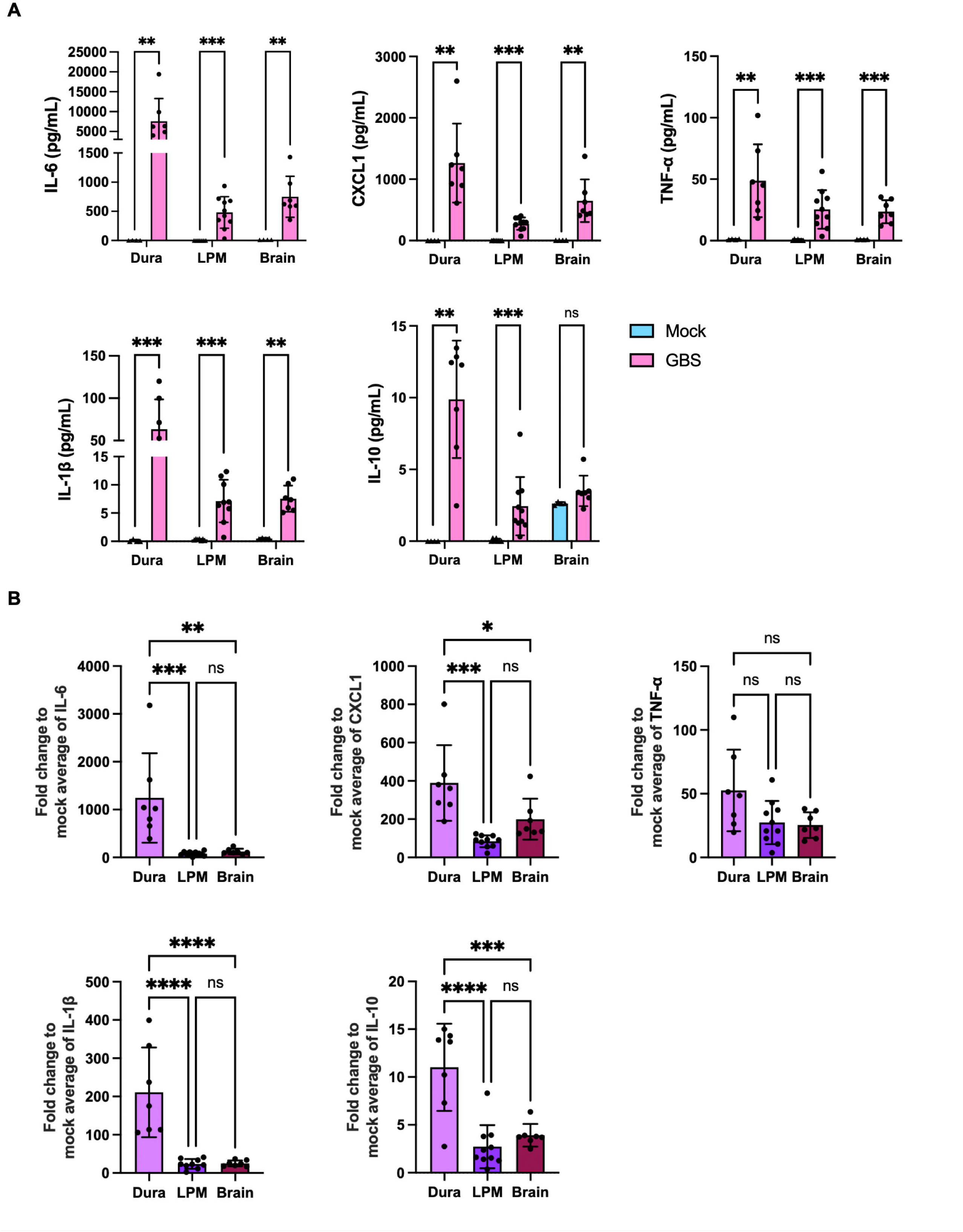
Proinflammatory response by meninges and brain upon GBS infection. **(A)** Multiplex analysis of proinflammatory cytokines and chemokines on mock- and GBS-infected dura, LPM, and brain. **(B)** Fold change of proinflammatory cytokines and chemokines between dura, LPM, and brain compared to average mock values. (n=4 dura mock, n=8 LPM mock, n=4 brain mock, n=7 dura GBS, n=10 LPM GBS, n=7 brain GBS) Statistics: (A): Mann-Whitney *U* test, (B): One-way ANOVA, * = p < 0.05, ** = p < 0.01, *** = p < 0.001, ns = not significant; mean and SD.

### L-BAMs play a minimal role in AB breakdown during GBS infection

BAMs are the predominant resident immune cell in the LPM though low numbers of neutrophils, dendritic cells, and T-/B-cells are also present in healthy mice (Goldmann et al., 2016; Van Hove et al., 2019). To investigate the immune cell contribution to proinflammatory cytokine production following GBS infection, we started by profiling the immune cell inventory of the LPM using flow cytometry (**Supp. Fig. 2B**). The overall percentage of L-BAMs, neutrophils, B-cells, and monocytes did not significantly change post-infection in LPM where L-BAMs accounted for 70-75% of the total CD45^+^ sorted cells (**Fig. 3A; Supp. Fig. 2B, C**). While the total number of L-BAMs (CD206^+^/Lyve1^+^) did not change after infection (**Fig. 3B, E**), they adopted a more polarized and elongated morphology post-infection (**Fig. 3D**) and Lyve1 expression was significantly decreased during GBS infection (**Fig. 3C, F**). Similar phenotypes have been reported in other inflammatory conditions and tissue resident macrophages (Drieu et al., 2022; Elfstrum et al., 2024; Johnson et al., 2007; Jordão et al., 2019; Lim et al., 2018; Wang et al., 2023). Other immune cell types were unaffected during infection except for a small increase in neutrophils from ∼2% to ∼4% of total live CD45^+^ cells (**Supp. Fig. 2C**). Taken together, these data show that L-BAMs are the primary immune cell population in the postnatal LPM and exhibit alterations in shape and a loss of Lyve1 expression following bacterial infection. Further, the overall number of L-BAMs does not change and there was only a small invasion of neutrophils at this early time point.

**Figure 3.**
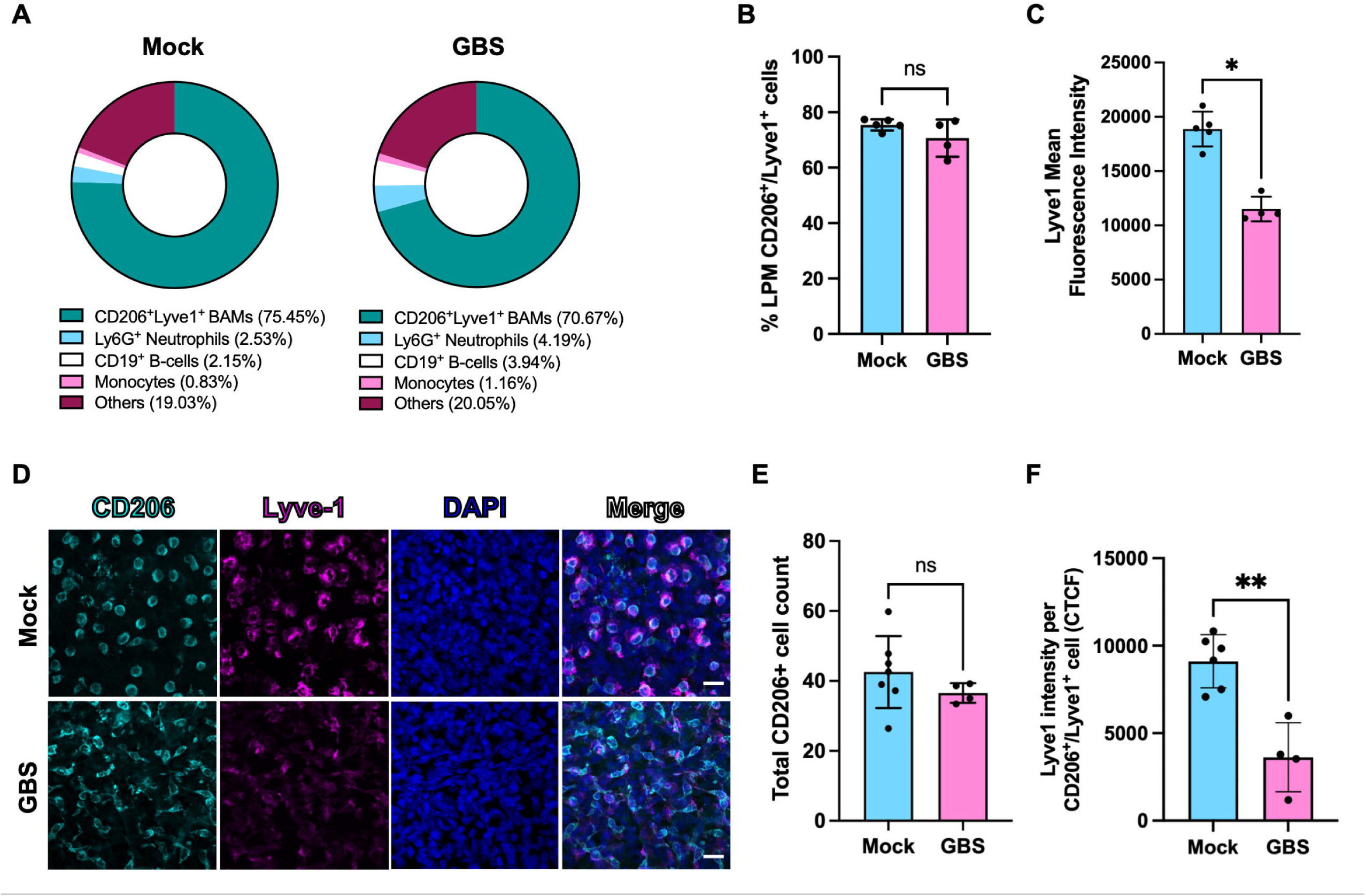
Immune response during GBS infection in neonatal LPM. **(A)** Flow cytometry analysis of % immune cells present in LPM whole mounts between mock- and GBS-infected group. **(B)** % LPM CD206^+^/Lyve1^+^ cells of live CD45^+^ immune cells and **(C)** Lyve1 mean fluorescence intensity of cells in mock- and GBS-infected group from (A). (n=5 mock, n=4 GBS for 3A-3C) **(D)** Representative images of L-BAMs using IF on LPM whole mounts. **(E)** Total CD206^+^ cell count and **(F)** Lyve1 intensity per CD206^+^/Lyve1^+^ cell (CTCF) from **(D)**. (n=7 mock, n=4 GBS for 3E and 3F) Statistics: Mann-Whitney *U* test, * = p < 0.05, ** = p < 0.01, ns = not significant; mean and SD. Scale bar = (D) 20µm.

We next examined the production of proinflammatory cytokines *Il6* and *Tnfa* by *Mrc1*^+^ (gene that encodes CD206) meningeal BAMs following GBS infection by performing RNAscope on whole head sections, which include the dura (attached to the calvarium that physically separates from the LPM and brain during tissue processing) and LPM attached to the brain (**Fig. 5A**). The RNAscope analysis showed increased *Il6* and *Tnfa* expression in *Mrc1*^+^ dural and L-BAMs in GBS-infected animals compared to mock-infected animals (**Fig. 4A, B; yellow arrowheads**). Next, we depleted L-BAMs by injecting clodronate intracerebroventricularly (ICV) 24 hours prior to GBS infection (P1) (**Fig. 4C**). We show that ICV clodronate injections successfully reduced CD206^+^ L-BAMs 24 hours post-injection (**Fig. 4C; Supp. Fig. 3A**) and that CD206^+^ dural BAMs are still present (**Supp. Fig. 3B**). We found that L-BAM depletion prior to GBS infection did not alter clinical outcomes, which include % weight change and bacterial CFU in blood, LPM, and brain (**Fig. 4D; Supp. Fig. 3C, D**). L-BAM depletion did not prevent the loss of AB integrity induced by GBS infections as measured by the transcranial dye assay (**Fig. 4E, F**). Next, we ran a multiplex cytokine assay on dissected LPM (which include fibroblasts, endothelial cells, and BAMs) and found that overall levels of IL-6 and TNF-α levels in LPM post-GBS infection did not change with L-BAM depletion (**Fig. 4G**). Of note, the level of CXCL1, which plays a key role in neutrophil recruitment, was increased in L-BAM depletion group (p-value = 0.0256), compared to vehicle group potentially due to the loss of anti-inflammatory macrophages that dampen the production of proinflammatory molecules by other cells or by compensatory recruitment of neutrophils (Chen et al., 2023). Collectively, our findings show that both dural and L-BAMs produce inflammatory cytokines following GBS infection and that even in the absence of L-BAMs, there is major inflammatory response to infection by the meninges that is sufficient to drive AB breakdown.

**Figure 4.**
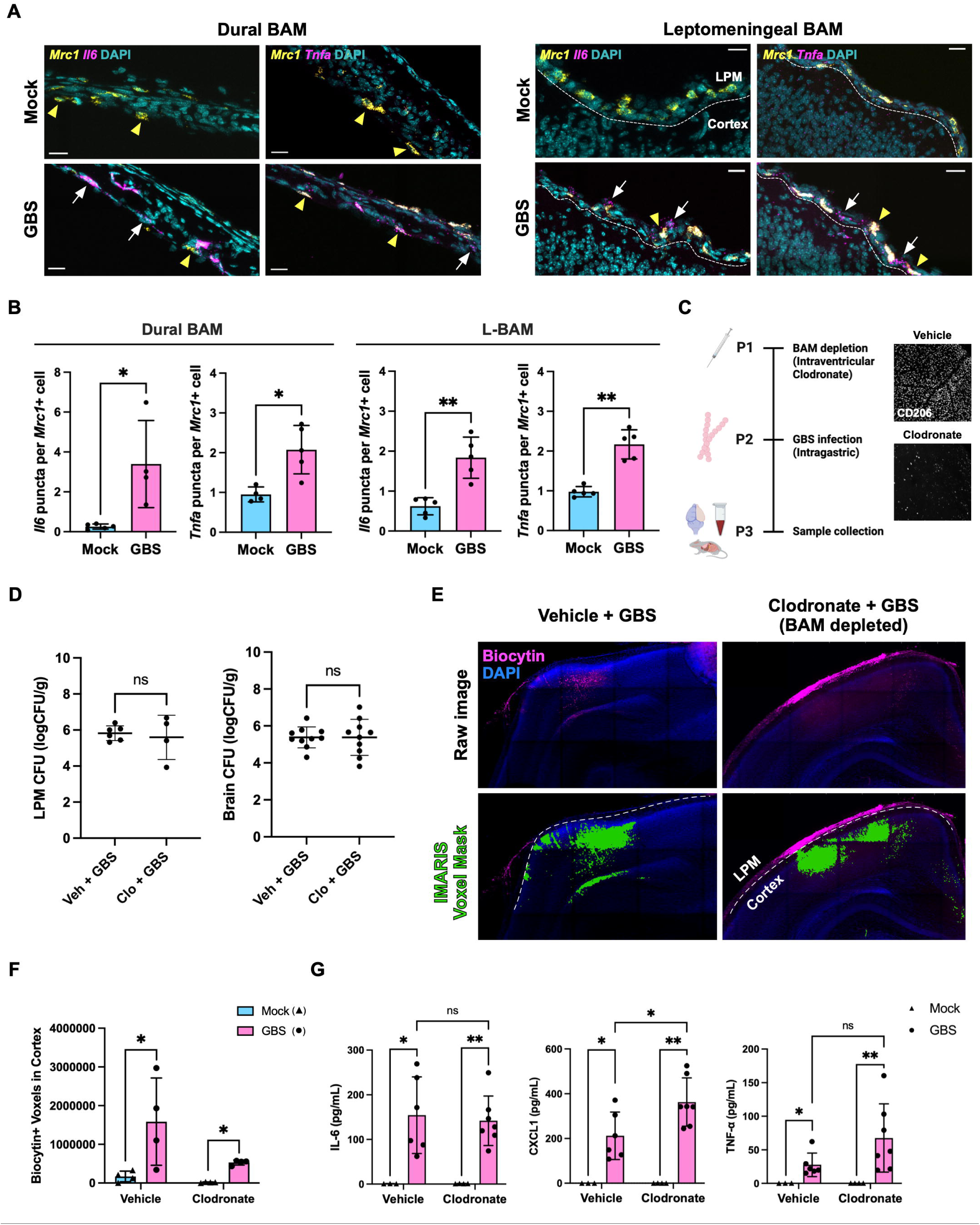
L-BAMs play a minimal role in AB breakdown during GBS infection. **(A)** Representative images of *Il6* and *Tnfa* expression in *Mrc1*^+^ dural- and L-BAMs from RNAscope-processed whole head sections. Yellow arrowheads indicate *Mrc1*^+^ cells and white arrows indicate *Mrc1*^-^ cells. **(B)** Quantification of *Il6* and *Tnfa* RNAscope punctas per *Mrc1*+ BAMs in dura and LPM. (n=5 mock, n=5 GBS) **(C)** Schematic of L-BAM depletion procedure in neonatal GBS infection model using clodronate. Insets show CD206^+^ BAMs in vehicle-vs clodronate-treated LPM. (Vehicle = empty liposome) **(D)** GBS CFU in LPM and brain in vehicle- and clodronate-treated mice. (n=6 mock LPM, n=4 GBS LPM, n=10 mock brain, n=11 GBS brain) **(E)** Qualitative images and **(F)** quantitative analysis of Biocytin-TMR intensity (IMARIS voxel mask) in the cortex using the transcranial AB integrity assay. (n=4 per group) **(G)** Multiplex assessment of IL-6, TNF-α, and CXCL1 levels in vehicle- and clodronate-treated mice in mock and GBS groups. (n=3 vehicle mock, n=6 vehicle GBS, n=4 clodronate mock, n=7 clodronate GBS) Statistics: (B), (D), (F): Mann-Whitney *U* test, (G): Two-way ANOVA, * = p < 0.05, ** = p < 0.01, ns = not significant; mean and SD. Scale bar = (A) 20µm.

**Figure 5.**
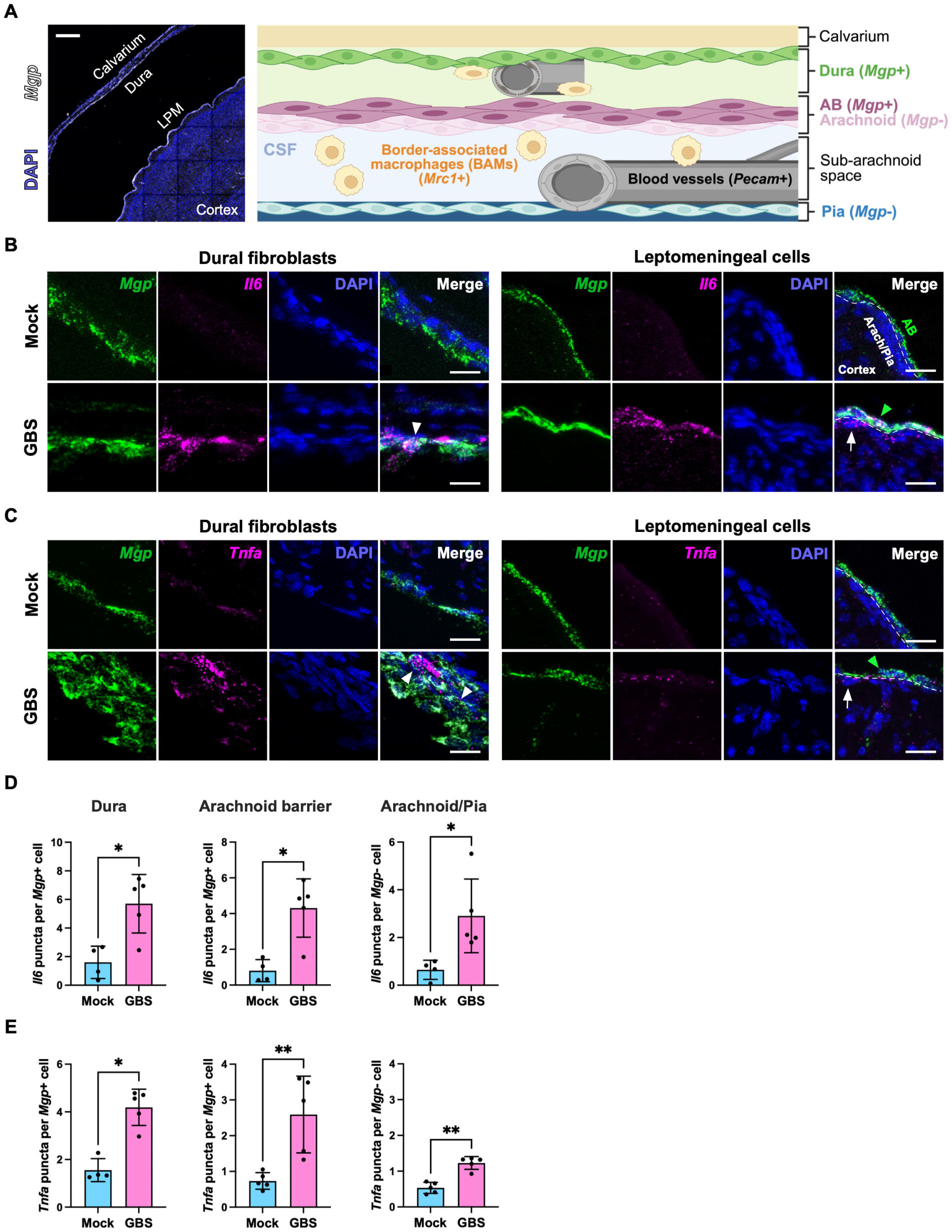
Production of proinflammatory molecules by meningeal fibroblasts during GBS infection. **(A)** Representative image of a whole head section (left) and graphical representation of meningeal cell types and gene expression (right). *Mrc1*^+^ (CD206^+^) dural- and L-BAMs, *Mgp*^+^ dural fibroblasts and AB cells, *Mgp*^-^ arachnoid/pial cells, and *Pecam*^+^ endothelial cells. **(B)** Representative images of *Il6* and **(C)** *Tnfa* transcripts in dural fibroblasts (left) and LPM (right) cells using RNAscope-processed whole head sections. Arrowheads indicate *Il6^+^* or *Tnfa*^+^ dural or AB cells (*Mgp*^+^); arrows indicate *Il6^+^* or *Tnfa*^+^ arachnoid/pial cells (fibroblasts, endothelia, L-BAMs) (*Mgp^-^*). **(D)** Quantification of *Il6* and **(E)** *Tnfa* RNAsocpe punctas in *Mgp*^+^ dural fibroblasts, *Mgp*^+^ AB cells, and *Mgp*^-^ arachnoid and pial cells. (n=4 mock, n=5 GBS for 5D and 5E) Statistics: Mann-Whitney *U* test, * = p < 0.05, ** = p < 0.01; mean and SD. Scale bars = (A) 200µm, (B), (C) 20µm.

### Meningeal fibroblasts can directly respond to bacteria and are a source of proinflammatory molecules during GBS infection

To investigate the non-immune cell sources of proinflammatory molecules post-GBS infection in the meninges, we performed RNAscope on whole head sections using probes for *Il6* and *Tnfa* along with the probe for Matrix gla protein (*Mgp*) and *Pecam*. *Mgp* marks dural fibroblasts and AB cells (DeSisto et al., 2020; Pietilä et al., 2023) and *Pecam* marks endothelial cells (**Fig. 5A; Supp. Fig. 3E**). The RNAscope analysis showed higher expression of *Il6* and *Tnfa* in *Mgp*^+^ dural fibroblasts, as well as in *Mgp*^+^ AB cells and *Mgp*^-^ arachnoid and pial cells in GBS-infected animals compared to mock infection (**Fig. 5B, C**). *Pecam*^+^ endothelial cells also showed increased expression of *Il6* and *Tnfa* post-infection (**Supp. Fig. 3E**). This analysis demonstrates that *Il6* and *Tnfa* expression are increased post-GBS infection by dural fibroblasts, AB cells, arachnoid/pial cells, and endothelial cells. (**Fig. 5B-E**).

To further examine the meningeal fibroblast-specific response to GBS infection, we exposed primary fibroblast cultures established from P3 dura or P3 LPM to GBS *in vitro,* performing quantitative assays for cytotoxicity, bacterial adhesion and invasion, and proinflammatory cytokine ELISA as previously described (Joyce et al., 2022a, 2024) (**Fig. 6A**). GBS adherence and intracellular invasion to host cells are two key steps to successful bacterial infection. Our data show that GBS adhere to and invade cultured LPM fibroblasts, but minimally with cultured dural fibroblasts (**Fig. 6B, C**). We next assessed proinflammatory cytokine production following GBS exposure by performing an ELISA analysis on cultured dural and LPM fibroblasts. The result showed that the levels of IL-6 and CXCL1 were significantly higher at 12h and 18h post-exposure for both LPM and dural fibroblasts (**Fig. 6D**). For TNF-α, LPM fibroblasts showed a modest increase in expression at 12h post-GBS exposure and dural fibroblasts did not show increases at any timepoint. Using lactate dehydrogenase (LDH) cytotoxicity assay, we confirmed the inflammatory responses were not due to cytotoxicity (**Supp. Fig. 4A**). These data collectively demonstrate that the meningeal fibroblasts are a source of proinflammatory cytokines upon GBS infection, and the robust inflammatory signaling observed *in vivo* is likely a collective response of meningeal fibroblasts to GBS as well as to molecules produced by other cell types in response to infection.

**Figure 6.**
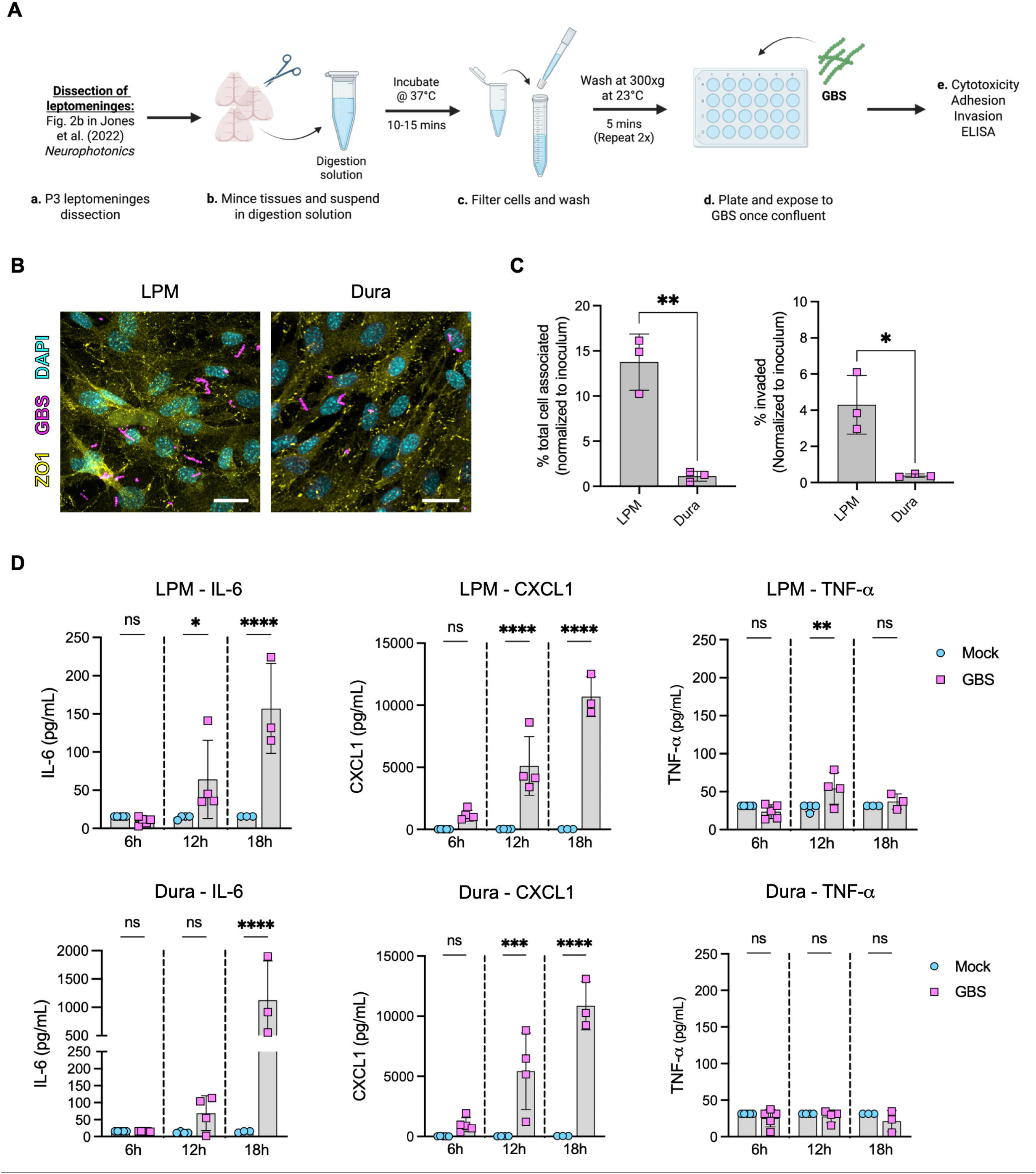
Meningeal fibroblast-specific response to GBS infection *in vitro*. **(A)** Schematic of primary meningeal fibroblast culture preparation and quantitative assays. **(B)** IF images of GBS on primary LPM and dural fibroblasts. **(C)** Adherence (% total cell associated) and invasion assays on LPM and dural fibroblasts. (n=3 per group) **(D)** Proinflammatory response by LPM (top) and dural (bottom) fibroblasts at 6, 12, 18 hours post-GBS exposure. (n=5 6hr, n=4 12hr, n=3 18hr) Statistics: Unpaired t-test, * = p < 0.05, ** = p < 0.01, *** = p < 0.001, **** = p < 0.0001, ns = not significant; mean and SD. Scale bars = (B) 20µm.

### Proinflammatory cytokine TNF-**α** is sufficient to induce AB breakdown

To determine if sustained inflammatory environment is sufficient to induce AB breakdown in the absence of GBS, we peripherally delivered TNF-α (300ng) to P2 pups and analyzed AB integrity at P5 (**Fig. 7A**). TNF-α is a well-established driver of barrier and junctional disruption, including in the human brain microvascular endothelium, human umbilical vein endothelium, and mouse intestinal epithelium (Al-Sadi, 2009; Capaldo and Nusrat, 2009; Gryka-Marton et al., 2025). While peripheral injection of TNF-α did not induce morbidity or significantly halt growth within the three days post-injection (**Fig. 7B**), the AB became significantly more permeable as indicated by an increased tracer leakage into the cortex of TNF-α-treated group (**Fig. 7C**). IF processing of LPM whole mounts showed that the expression of junctional Cldn11 was not different between the two groups (**Fig. 7D, E**), but the total length of Cldn11^+^ junctions was significantly lower in the TNF-α-treated LPM (**Fig. 7F**), like what was observed in Cldn11 junctions in GBS-infected animals. Taken together, these results demonstrate that the proinflammatory cytokine TNF-α is sufficient to disrupt the neonatal AB and serves as a potential factor underlying aberrant distribution of junctional proteins and barrier leakage in bacterial meningitis.

**Figure 7.**
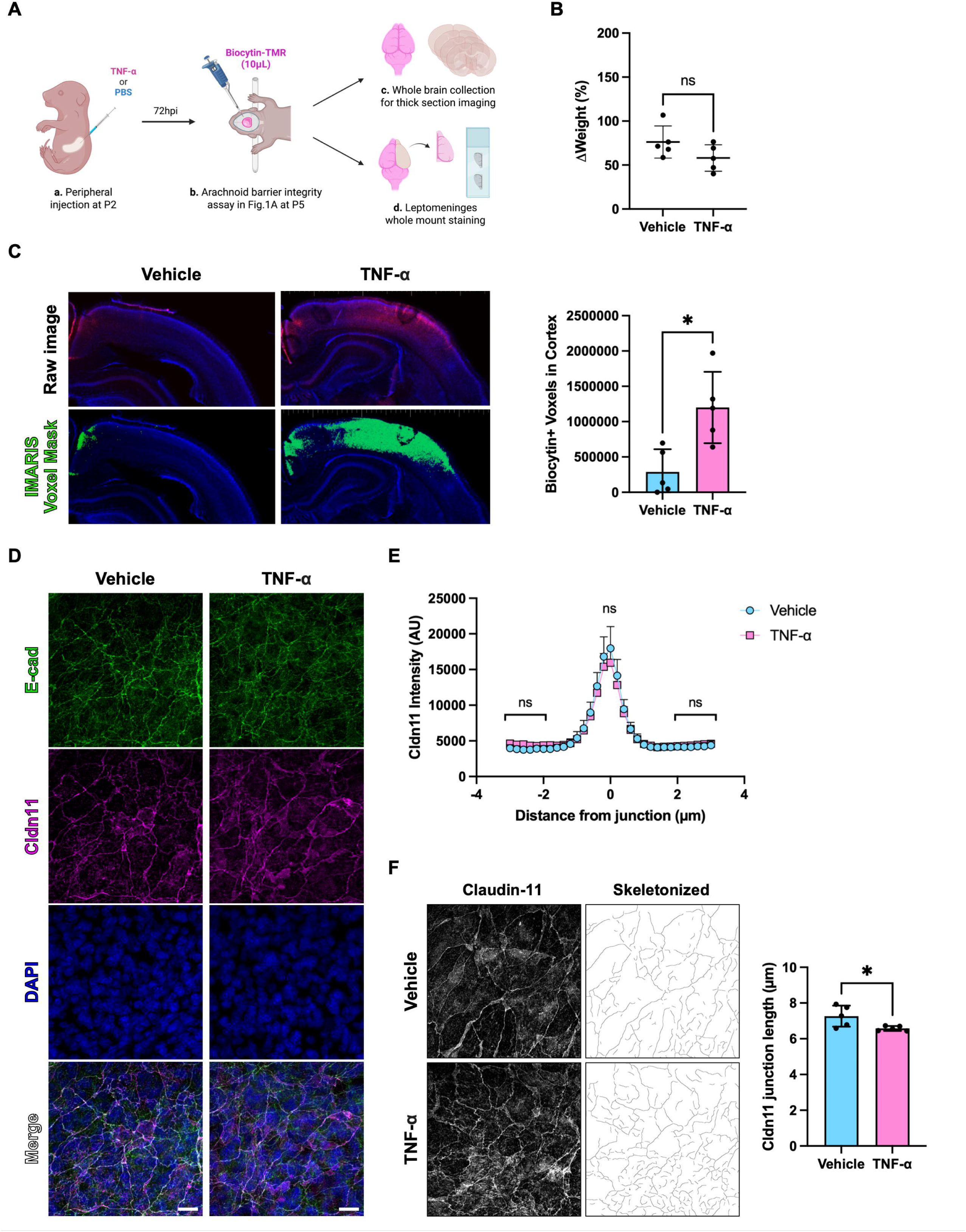
Proinflammatory cytokine TNF-α is sufficient to induce AB breakdown. **(A)** Schematic of testing the effect of peripheral TNF-α administration on neonatal AB integrity. **(B)** % weight change in vehicle (PBS)- and TNF-α-treated groups between P2 and P5. **(C)** Qualitative images and quantitative analysis of transcranial AB integrity assay post-peripheral TNF-α administration. (500µm coronal brain sections) **(D)** Representative images of E-cad^+^ and Cldn11^+^ AB junctions in LPM whole mounts post-TNF-α exposure. **(E)** Line intensity analysis of Cldn11^+^ junctions. **(F)** Ridge Detection analysis of Cldn11^+^ junctions. (n=5 per group in 7B, 7C, 7E, 7F) Statistics: Mann-Whitney *U* test, * = p < 0.05, ns = not significant; mean and SD. Scale bars = (D) 20µm.

## DISCUSSION

Neonatal meningitis is a leading cause of morbidity and long-term neurological dysfunction. However, the pathophysiology remains only partially understood with large gaps in our understanding of how the AB and meningeal fibroblasts respond to infection. Here, we demonstrate for the first time that bacterial meningitis causes loss of AB integrity and show that meningeal fibroblasts contribute to proinflammatory cytokine production that is disruptive to AB integrity. Using a GBS neonatal meningitis model, we document increased permeability across the AB and disruption of Cldn11^+^ tight junction organization in AB cells. We also found a significant upregulation of proinflammatory molecules in the meningeal space and the brain, including IL-6, CXCL1 and TNF-α, and activation of L-BAMs. Surprisingly, we found that clodronate-mediated L-BAM depletion prior to infection did not improve GBS infection-induced AB breakdown or significantly reduce the levels of proinflammatory cytokines in the LPM post-infection. *In situ* and cell culture assays revealed that meningeal fibroblasts, BAMs, and endothelial cells are involved in the production of proinflammatory molecules in response to GBS and highlight meningeal fibroblasts as an important source. Lastly, we show that peripheral injection of TNF-α, a barrier disruptive proinflammatory cytokine upregulated in a range of CNS infections and diseases, is sufficient to cause AB breakdown and mislocalization of tight junction protein Cldn11.

The AB is a critical component of the B-CSFB that physically separates the CSF-filled subarachnoid space from permeable vasculature of the overlying dura. The AB controls the movement of peripheral molecules into the CSF and regulates CSF composition through expression of efflux transporters and solute carrier (SLC) transporters (Uchida et al., 2020; Yasuda et al., 2013; Zhang et al., 2018). However, how the AB responds to insults and the consequences of its breakdown are largely unknown. Here, we report for the first time that bacterial meningitis infection in neonates can induce AB breakdown. With a recent surge in studies of the AB, including the descriptions of its developmental maturation and advances in methods to visualize and functionally test the AB integrity (Derk et al., 2023; Mapunda et al., 2023b; Smyth et al., 2024c; Zhang et al., 2022), the field is now poised to advance studies on the consequences of AB breakdown in brain health and function. A recent work showed that in mice missing an innate-like T-cell population called MAIT cells that reside in the meninges, the AB lost its barrier integrity and the mice developed a pronounced microgliosis and impaired hippocampal-mediated learning (Zhang et al., 2022). Although the GBS infection model used here induced morbidity, modifications to this model in the future will allow us to investigate the consequences of AB breakdown and if the AB is capable of being repaired following resolution of the infection and local inflammation.

Our results identify the production of inflammatory cytokines in the meninges post-GBS infection and we further show that systemic TNF-α exposure, even in the absence of GBS, is sufficient to induce AB breakdown. We found that IL-6 and CXCL1 were both significantly upregulated upon GBS infection. These molecules are known to function as chemoattractant for neutrophils and stimulate the expression of other cytokines (Karimbakhsh et al., 2026; Michael et al., 2020; Nie et al., 2025; Roy et al., 2012), suggesting roles for these cytokines analyzed here. Systemic and local TNF-α is likely a major driver of AB breakdown during infection because it has been shown to induce BBB and tight junction breakdown in various neurological diseases and models (Al-Sadi, 2009; Capaldo and Nusrat, 2009; Gryka-Marton et al., 2025). *In vitro* studies in epithelial and endothelial cells have reported a significant decrease in transepithelial/transendothelial resistance (TEER) upon TNF-α exposure (Al-Sadi, 2009) and have shown that TNF-α induces disruption of tight junctions proteins ZO-1, occludin and claudin-5 (Al-Sadi, 2009; Ma et al., 2004; McKenzie and Ridley, 2007). Thus, it is possible that in other neurological diseases and infections where levels of TNF-α spike, the AB will be compromised.

A main goal of our studies is to understand the functions of meningeal fibroblasts and here we report that meningeal fibroblasts are an important source of proinflammatory cytokines during GBS infection. Our findings are consistent with other studies and underscore the idea that CNS fibroblasts play an important role in disease and infection progression. For example, following viral infection, perivascular fibroblasts produce CCR7 to promote activation of antiviral CD8^+^ T cells (Cupovic et al., 2016), while a different study showed that PDGFRβ+ perivascular fibroblasts, but not microglia or astrocytes, are the initial source of CCL2 following lipopolysaccharide administration (Duan et al., 2018). A recent study identified substantial brain fibroblast-immune cell coordination that occurs following brain injury, and the important role that brain fibroblasts play in modulating inflammatory response and limiting tissue loss post-injury (Ewing-Crystal et al., 2025). Thus, our characterization of meningeal fibroblast response to GBS infection contributes to the growing literature describing the critical roles these cells play in infection and CNS diseases.

GBS is known to induce pathology by directly binding to host cells. We show that GBS can attach to and invade meningeal fibroblasts, consistent with a previous study showing GBS interaction with leptomeningeal-like human meningioma cells (Alkuwaity et al., 2012). Notably, while we observed no LDH-mediated cytotoxicity, GBS invasion led to significant cytokine production *in vitro*, a response not previously characterized in this cell type. Our data support that direct GBS-meningeal fibroblast interaction is a potent driver of proinflammatory cytokine production *in vivo* and set up the prediction that GBS interaction with AB cells induces cellular and molecular changes that contribute to AB breakdown and dysfunction. Meningitis-inducing bacteria, such as *Neisseria meningitidis,* have been shown to trigger proinflammatory responses in the human meningioma cells (Hardy et al., 2000; Christodoulides et al., 2002). Furthermore, single-cell profiling of the meninges in a postnatal *E. coli* meningitis model identified significant gene expression changes in the AB cell cluster, including downregulation of *Slc* transporter genes that underlie AB function at the B-CSFB (Wang et al., 2023). Beyond fibroblasts, GBS utilizes various surface proteins and toxins to interact with brain endothelium, promoting bacterial uptake and increasing barrier permeability (Mu et al., 2014; Coureuil et al., 2017; Joyce et al., 2024; Yang et al., 2023; Deng et al., 2019). Specifically, it has been demonstrated that a GBS surface protein BspC interacts with vimentin expressed by brain endothelial cells—a protein also highly expressed in arachnoid fibroblasts and by E-cad^+^ AB cells (Pietilä et al., 2023a; Shah et al., 2023)—to drive the pathogenesis of GBS meningitis (Deng et al., 2019). These interactions could disrupt barrier integrity, potentially through Snail1-mediated disruption of tight junction proteins shown in brain endothelium (Kim et al., 2015), and ultimately facilitate bacterial brain entry and promote a proinflammatory signaling cascade.

The focus of our study was the response of the AB and meningeal cells in neonates to bacterial infection. However, understanding why a significant percentage of newborns develop long-term neurological sequelae is particularly important (Furuta et al., 2022). It is possible that chronic changes to meningeal cells (fibroblast, vascular, and immune populations) and CNS barriers contribute to prolonged neurological symptoms and are therefore a potential target for therapeutic intervention. One important question to address in the future is whether the early infection could “prime” meningeal immunity that is being established during the postnatal period (Van Hove et al., 2019; Kim et al., 2023a; Barron et al., 2024; Walker et al., 2025) and whether this is beneficial or detrimental to prolonged neurological symptoms. Another is whether the AB can repair itself after the initial insult and if so, what mechanisms improve or prohibit this healing to occur. Further, it will be important to address if functional restoration of AB integrity is possible, if there is resolution of fibroblast cytokine production once the initial bacterial infection has been cleared, and if the dosage of GBS, timing of infection, or genetics of the host impact the outcome. Elucidating these mechanisms in future studies will be vital for developing novel therapies which promote health of the CNS through improved structural and functional integrity of the meningeal CNS barriers.

## MATERIALS AND METHODS

### Animals

Mice used for experiments here were housed in specific-pathogen-free facilities approved by the AALAC and were handled in accordance with protocols approved by the University of Colorado Anschutz Medical Campus IACUC committee. Mouse used in this study were wildtype mice on the C57Bl/6J background (Jackson: 000664). Pregnant females were checked daily for litters and the day of birth was considered postnatal day 0.

### Bacterial culture

GBS COH1 strain (Kuypers et al., 1989) was grown statically at 37°C in Todd-Hewitt Broth (THB) as previously described (Joyce et al., 2024).

### Murine neonatal model of GBS meningitis

Postnatal day (P)2 male and female C57Bl/6J mice were injected via intragastric injection with 2×10^6^ CFU of COH1 GBS or phosphate-buffered saline (PBS) as previously described (Joyce et al., 2024). Mice were monitored for weight loss, righting reflex, exhaustion levels, and dehydration over 18-24 hours (**Supp. Fig. 1B**). At 24 hours post-infection (hpi) or moribund state, mice were euthanized, and the blood and tissues were harvested, homogenized, and serially diluted on GBS CHROM agar plates to determine bacterial CFU (**Supp. Fig. 1C**).

### Transcranial arachnoid barrier integrity assay (ABIA)

This assay was adapted from a previously described transcranial SR101 assay (Zhang et al., 2022). At the time of tissue harvest, the animals were anesthetized via intraperitoneal injection of sodium pentobarbital (40mg/kg). Upon checking pedal reflex, their heads were rested on a flat post and calvarium was carefully exposed using sterile dissection scissors. The surface of calvarium was dried and sterilized with 70% ethanol and a 10 µL of fluorescent dye Biocytin-TMR (0.1% in saline; 870Da; Invitrogen, #T12921) was applied to the sagittal suture between bregma and lambda for 10 minutes. 10 µL of 0.1% Biocytin was reapplied at 5 minutes to prevent drying. Animals were placed on a heating pad throughout the entire procedure and humanely sacrificed by isoflurane overdose and decapitation at the end of the experiment. Key variables allowing for this experiment are 1) the calvarium is porous and 2) the dura (top layer of meninges) is devoid of a barrier layer; therefore, the dye permeates calvarium and dura layers during this assay and is impeded from entering the brain by an intact AB layer.

For quantitative analysis of Biocytin dye permeability across the AB in **Fig. 1**, the LPM and a piece of superficial cortex directly beneath the region of dye application were excised using sharp forceps and homogenized using a dounce homogenizer. The samples were processed and measured on a plate reader for: 1) Biocytin fluorescence intensity reading and 2) BCA Protein Assay (ThermoFisher, #23225) for sample normalization. For quantitative analysis of Biocytin dye in **Fig. 4** and **Fig. 7**, whole brains were collected, fixed with 4% PFA, and sectioned on a vibratome at 500µm thickness. The amount of Biocytin dye was quantified as total voxels in cortex on IMARIS.

### Clodronate-mediated macrophage depletion

P1 C57Bl/6J mice (24 hours prior to infection) were cryo-anesthetized for 3 minutes. Following cessation of movement, clodronate-containing liposomes (3µL of 5mg/ml stock solution; Encapsula Nano Sciences, #CLD-8914) or vehicle liposomes were injected into the right lateral ventricle using a sterile 10µL Hamilton syringe (Hamilton, 7653-01) with a 32-gauge needle (Hamilton, 7803-04). The injection site was located around two-fifths of the distance between lambda and right eye (Kim et al., 2013). Following injection, mice were placed on a heating pad until active and returned to their original cage with dam.

### Dissociation of LPM and staining for flow cytometric analysis

At the time of harvest (P3; 18-24hpi), animals were euthanized then perfused with sterile HBSS with Ca^2+^/Mg^2+^ (Invitrogen, #14025092). LPM tissues were harvested following our previous protocol (Jones et al., 2022). For single cell suspensions, LPM tissues were pooled from 3 mice per sample and digested in 1.5 mL of digestion solution consisting of HBSS, 2% w/v bovine serum albumin, 1% w/v glucose, 5 mg/mL Type II collagenase (Worthington, #NC9870009) and 2% DNase for 15 minutes at 37°C. The tissues were then triturated until well dispersed in the digestion solution. Liberation of the cells from the tissues into a single cell suspension was confirmed by viewing samples of the solution under a microscope, with additional incubation time at 37°C if needed to completely disaggregate the cells.

Single cell suspensions were first stained with eBioscience Fixable Viability Dye eFluor 506 (Catalog # 65-0866-18) in PBS for 30 minutes at room temperature. Cells were stained with the following anti-mouse surface antibodies in MACS buffer for 30 minutes at room temperature: from BioLegend—F4/80-BV785 (clone BM8; Catalog #123141), Ultra-LEAF Purified CD16/32 (clone 93; Catalog # 101330]; from BDBiosciences—CD19-PE-Cy5 (clone 1D3; Catalog # 558079), CD45-BUV395 (clone 30-F11; Catalog # 564279), IA/IE-AF700 (clone M5/114.15.2; Catalog # 748845), Ly6C-PercP (clone AL-21; Catalog # 562727); from Miltenyi Biotec—CD11b-APCVio 770 (clone M1/70; Catalog # 11-0112-41); from ThermoFisher Invitrogen—CD206-PE-Cy7 (clone 1A8-Ly6g; Catalog # 17-9668-82), Ly6G-APC (clone 1A8-Ly6g; Catalog # 17-9668-82), Lyve1-PE (clone 1A8-Ly6g; Catalog # 17-9668-82), P2RY12-AF488 (clone 1A8-Ly6g; Catalog # 17-9668-82). After surface antibody staining, the cells were fixed for 30 minutes at room temperature using the FoxP3 fixation/permeabilization kit (ThermoFisher Scientific, Catalog # 00-5523-00). Stained cells were analyzed on Cytek Aurora using the SpectroFlo software (v9). Data were analyzed with BD FlowJo software v 10.10.0. Gating strategy can be found in **Supp. Fig. 2B**.

### Multiplex cytokine array

Cytokines were measured using a multiplex cytokine array (V-PLEX Plus Proinflammatory Panel 1 Mouse Kit; #K0082534) according to the manufacturer’s instructions. Briefly, the tissues (dura, LPM, brain, liver) were homogenized using RIPA Lysis and Extraction Buffer (ThermoFisher, #PI89900) containing 2X Protease Inhibitor Cocktail (Roche, #4693116001) and a 50µL of aliquot was taken for BCA Protein Assay. Remaining samples were stored at -80°C until use, which were thawed and spun at 2,000 x g for 3 minutes. Pre-coated V-PLEX plates were washed using an automated plate washer (BioTek ELX5012), 50µL of Calibrators or diluted samples were added, and plates were incubated for 2 hr at room temperature on a Compact Digital Microplate shaker (ThermoFisher) at 600 rpm. The plates were washed and 25µL of diluted detection antibodies were added and incubated for 2 hr at room temperature. After washing, 2× Read Buffer (MesoScale Discovery) was added and plates were immediately read on a MesoQuickPlex SQ120 electro-chemiluminescent plate reader. Data were analyzed using Workbench software (MesoScale Discovery).

### Primary meningeal fibroblast culture

Primary dural and LPM fibroblasts were collected from P3 C57Bl/6J mice. Following euthanasia, the dura and LPM were carefully dissected in cold, sterile HBSS from calvarium and brains, respectively. The LPM or dura tissues were pooled from 3 mice per sample, and the tissues were dissociated using the dissociation procedure for flow cytometric analysis described above and visualized in **Fig. 6A**. After successful dissociation and single cell suspension, cells were seeded on Matrigel-coated well plates and incubated in DMEM/F-12 medium (Gibco, #11330032) containing 10% fetal bovine serum and 1% Penicillin-Streptomycin at 37°C.

### GBS adherence, invasion, and cytotoxicity *in vitro* assays

Adherence and invasion assays were performed as previously described (Joyce et al., 2022b) on confluent cells (∼2×10^5^ cells) in a 24-well tissue culture treated plate (Corning). Briefly, for the adherence assay, cells were infected in technical triplicate at a multiplicity of infection (MOI) of ∼1 for 30 minutes. Cells were gently washed 4 times with PBS, trypsinized, and lysed for plating bacterial counts. For invasion assays, cells were infected in triplicate at an MOI of ∼1 for 2 hr, media was replaced with fresh media supplemented with 100 µg of gentamycin and 5 µg of penicillin and incubated for another 2 hr. Cells were gently washed once, trypsinized, and lysed for plating bacterial counts.

Cytotoxicity was measured via lactate dehydrogenase (LDH) release assay (CyQUANT, #C20300) per manufacturer protocols. Media was replaced with DMEM (no phenol red or serum) and infected at an MOI of ∼50 or treated with 0.1% Triton (maximum LDH release) for 6-, 12-, and 18-hr time points. Whole supernatants were collected and spun down to remove debris, then plated in duplicates for measuring LDH release. Cytotoxicity was determined by considering the spontaneous release (UI) and the maximum release (Triton).

### Enzyme-linked immunosorbent assay

Supernatants were collected from uninfected and infected LPM and dura cultures at 6, 12, and 18 hrs post-GBS exposure. Murine CXCL-1, IL-6, and TNF-α in supernatants were detected by ELISA according to the manufacturer’s instructions (R&D systems).

### Immunocytochemistry

Upon reaching desired confluency on a round glass coverslip (12mm) coated with Matrigel, cells were incubated with GBS for 1hr at an MOI of ∼5, then washed twice with 1X PBS and fixed with 4% PFA for 10 minutes at room temperature. After washing twice with 1X PBS, cells were incubated in blocking buffer consisting of 10% lamb serum and 0.1% Triton-X in 1X PBS for 1 hr at room temperature. Following this, the blocking buffer was replaced with primary solution made up of primary antibodies, 2% lamb serum, and 0.1% Triton-X in 1X PBS and the cells were incubated on a shaker overnight at 4°C. On the following day, cells were washed twice with 1X PBS and incubated in secondary solution made up of secondary antibodies, 2% lamb serum, and 0.1% Triton-X in 1X PBS for 1 hr at room temperature. Lastly, cells were washed twice with 1X PBS and carefully mounted with Fluoromount-G Mounting Medium (Invitrogen). Primary antibodies (1:200 dilution) used here include: rabbit anti-GBS (GeneTex, #GTX40901) and mouse anti-ZO-1 (Invitrogen, #33-910-0). Appropriate combination of AlexaFluor-conjugated secondary antibodies (1:500; Invitrogen) and DAPI (1:1000; Invitrogen) were used.

### TNF-**α** treatment

Mouse TNF-α Recombinant Protein (PeproTech, #315-01A) was reconstituted in sterile PBS before use and 300ng was injected intraperitoneally at P2. Mice were monitored for overall health, such as weight, and the whole brains were collected at P5 following transcranial dye assay or LPM whole mounts were collected for junctional analyses.

### Immunostaining and RNAscope

Procedures for LPM whole mount dissection, fixation and immunostaining are detailed at length in our protocol paper (Jones et al., 2022). For cryosections of whole head used for RNAscope and brains prepared for vibratome sectioning, mice were euthanized by injecting a fatal dose of pentobarbital followed by transcardiac perfusion with 1X PBS followed by 4% paraformaldehyde (PFA). Heads (skin and lower jaw removed) were post-fixed in 4% PFA for 24 hours, then in 20% sucrose solution for 24 hours at 4°C. Prior to cryosectioning (12µm), the heads were embedded in Optimal Cutting Temperature (OCT) compound and flash-frozen in pre-cooled isopentane. For thick sections (100-500µm), brains were post-fixed in 4% PFA for 24h and stored in PBS until embedded in 3% agarose gel (1.5% low-melt agarose and 1.5% agarose) prior to vibratome sectioning. Incubation in the following primary antibodies (1:200 dilution) was conducted overnight at 4°C in appropriate solution (as described in Jones et al., 2022): mouse IgG2a anti-E-cadherin (BD Transduction Laboratories, #610181), rabbit anti-Claudin 11 (ThermoFisher, #PIPA568608), goat anti-CD206 (R&D Systems, #AF2535), rabbit anti-Lyve-1 (Abcam, #NC9819988), rat anti-PECAM (BD Biosciences, #BDB550274), mouse anti-ZO-1 (Invitrogen, #33-910-0). Following incubation with primary antibodies, tissue sections were incubated for 60 min at room temperature with appropriate AlexaFluor-conjugated secondary antibodies (1:500; Invitrogen) and DAPI (1:1000; Invitrogen). For RNAscope *in situ* hybridization, the tissues were processed following the manufacturer’s protocol (ACD Bio) and the following probes used on cryosections were: *Il6* (315891)*, Mgp* (463381)*, Mrc1* (437511)*, Pdgfra* (411361)*, Pecam* (316721)*, Tnfa* (311081).

### Image analysis

Images were obtained on Zeiss LSM 780 and 900 confocal microscope and Zen Blue software (Zeiss). For junctional line analysis used to measure the protein expression (**Supp. Fig. 1D**), a 10µm line was drawn perpendicularly across 10 randomly selected junctions per image and the “junctional” (peak at +0-0.45µm) and “non-junctional” (between +2-3µm) fluorescence intensities were measured using Plot Profile in FIJI. For ridge analysis used to assess the length or “continuity” of junctions (**Supp. Fig. 1E**), the Ridge Detection plugin in FIJI was used. To quantify the gene expression in RNAscope-processed tissues, images were analyzed as # of punctas per cells on IMARIS (Oxford Instruments). The number of *Il6* and *Tnfa* puncta was quantified per *Mrc1*^+^, *Mgp*^+^, *Mgp*^-^ cells (all DAPI^+^) in dura and leptomeninges. For quantification of AB integrity assay, total Biocytin+ voxels in the cortex were quantified by creating Biocytin+ surfaces on IMARIS.

## Statistical analysis

All statistics were performed using GraphPad Prism software V10. All replicates and statistical analyses are provided in figure legends.

## Acknowledgements

We thank the members of Drs. Julie Siegenthaler, Kelly Doran, and Stephanie Bonney’s labs for constructive feedback on the manuscript. We thank the members of the Human Immune Monitoring Resource (HIMSR, RRID:SCR_021985) within the University of Colorado Cancer Center (P30CA0434) for their help with multiplex cytokine analyses. We thank the ImmunoMicro Flow Cytometry Core Facility at the University of Colorado Anschutz for help with flow cytometry experiments. All graphical schematics were made in BioRender. This work was supported by R01 NS098273 to J.A.S. and R01 NS116716 to K.S.D. from NIH/NINDS.

## Author contributions

Conceptualization: S.K., L.J., K.S.D., J.A.S. Methodology: S.K., L.J., A.B., B.S., B.P., J.D., K.S.D., J.A.S. Validation: S.K., L.J., A.B., B.S., B.P., K.S.D., J.A.S. Formal analysis: S.K., L.J., A.B., B.S., B.P. Investigation: S.K., L.J., A.B., B.S., B.P. Writing—original draft: S.K. Editing and reviewing: S.K., L.J., A.B., B.S., B.P., J.D., K.S.D., J.A.S. Visualization: S.K., L.J., A.B., B.S., B.P. Project administration: K.S.D., J.A.S.

