## SupplementaryFigures for "Meningeal inflammation and arachnoid barrier breakdown in a mouse model of neonatal bacterial meningitis"

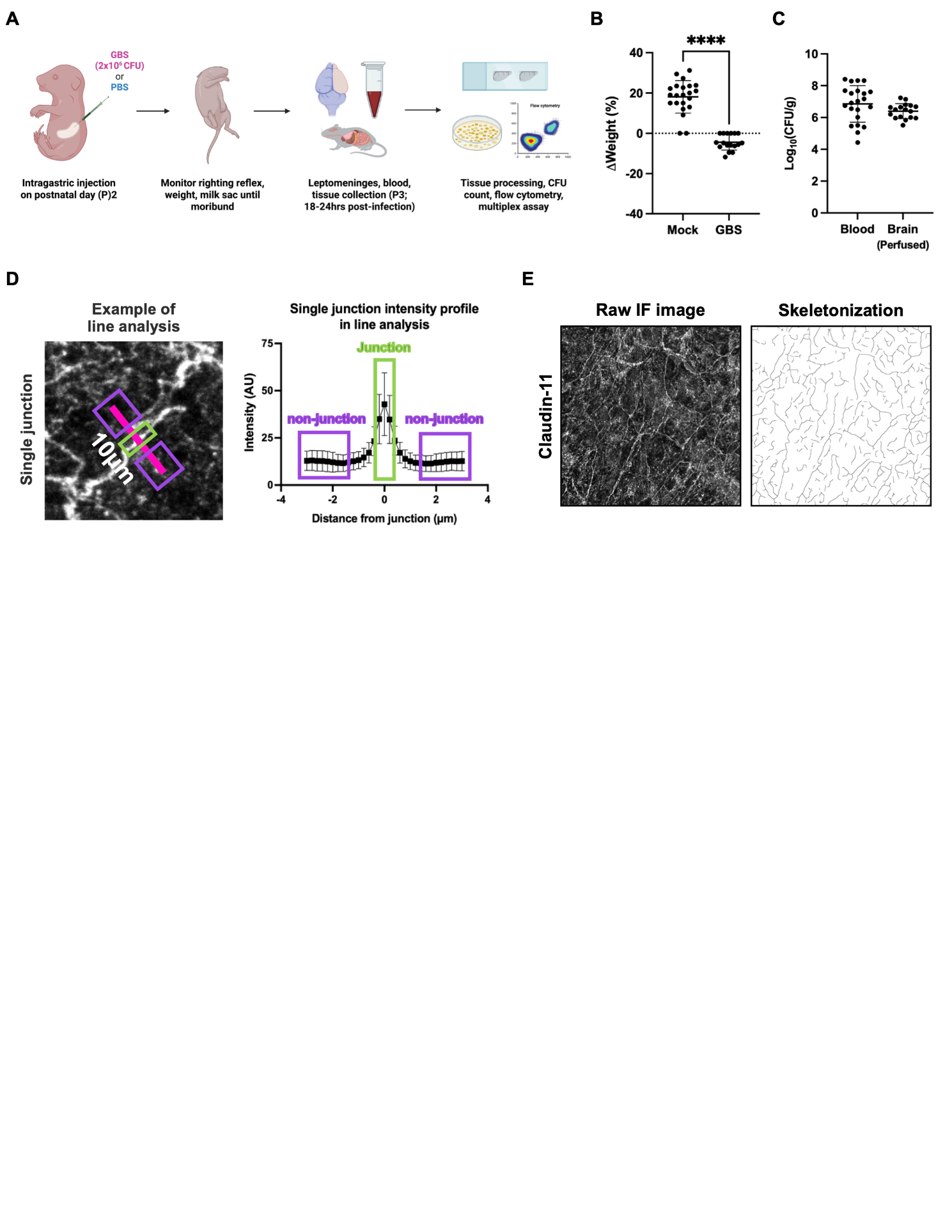

**Supplementary Figure 1. (A)** Graphical representation of bacterial meningitis model in neonatal mice using GBS. **(B)** % weight change between P2 and P3 in mock- and GBS-infected groups. (n=22 mock, n=17 GBS) **(C)** Blood and brain (perfused) CFU in the neonatal meningitis model. (n=22 blood, n=17 brain) **(D)** Schematic of single junction fluorescence intensity analysis using Plot Profile in FIJI ImageJ. A 10µm line was drawn perpendicularly to each junction and the intensity of the junction was measured at the peak (0-0.45µm) of the plot and non-junctional intensity at 2-3µm away from the peak. **(E)** Schematic of total junction length analysis using Ridge Detection plugin in FIJI ImageJ. Statistics: Mann-Whitney *U* test, **** = p < 0.0001; mean and SD.

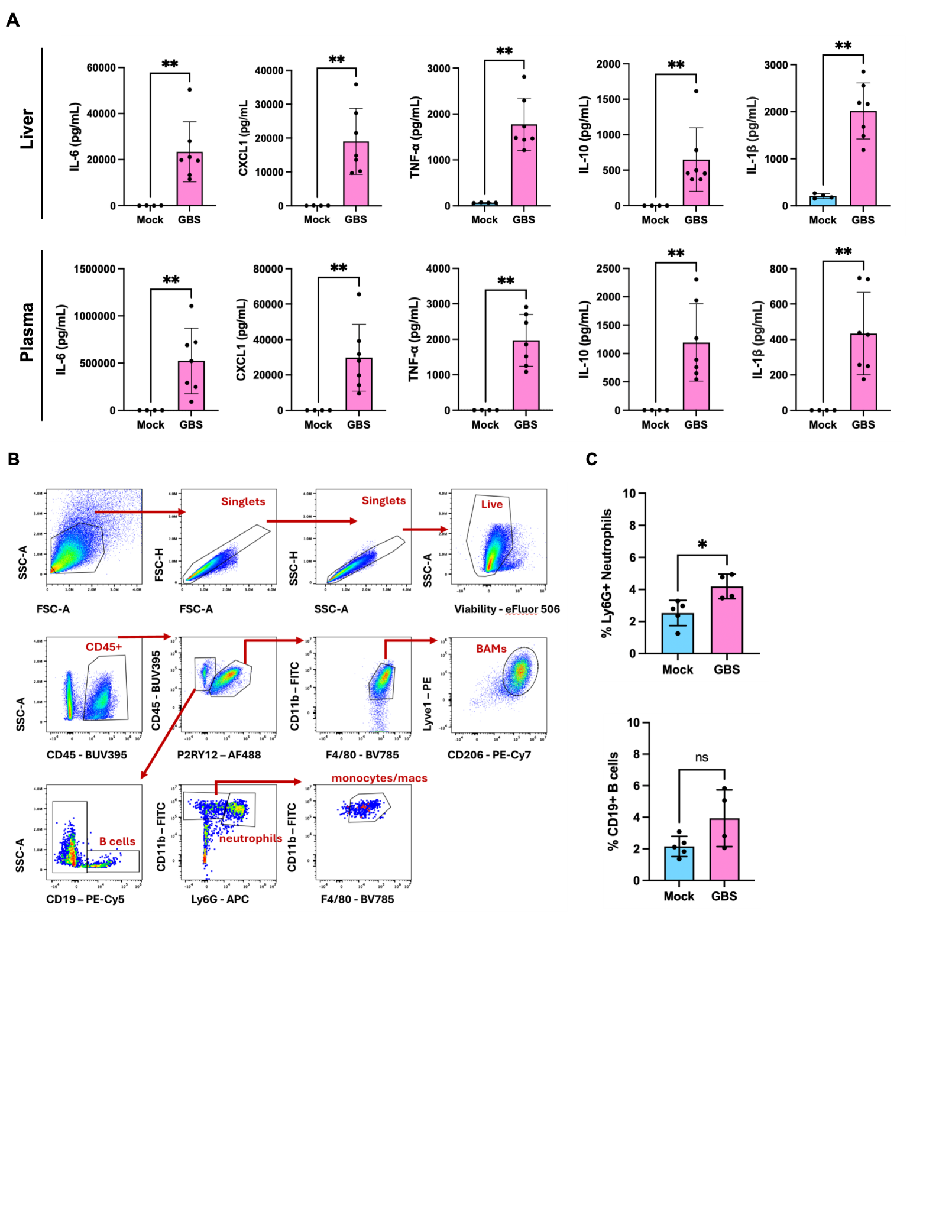

**Supplementary Figure 2. (A)** Multiplex quantification of proinflammatory molecules in liver and plasma post-GBS infection. (n=4 mock, n=7 GBS) **(B)** Gating strategy for flow cytometric analysis of LPM immune cells in Fig. 2(A). **(C)** Quantification of % Ly6G^+^ neutrophils (top) and % CD19^+^ B-cells (bottom) between mock- and GBS-infected groups of live CD45^+^ LPM immune cells. (n=5 mock, n=4 GBS) Statistics: Mann-Whitney *U* test, * = p < 0.05, ** = p < 0.01, ns = not significant (p-value > 0.05); mean and SD.

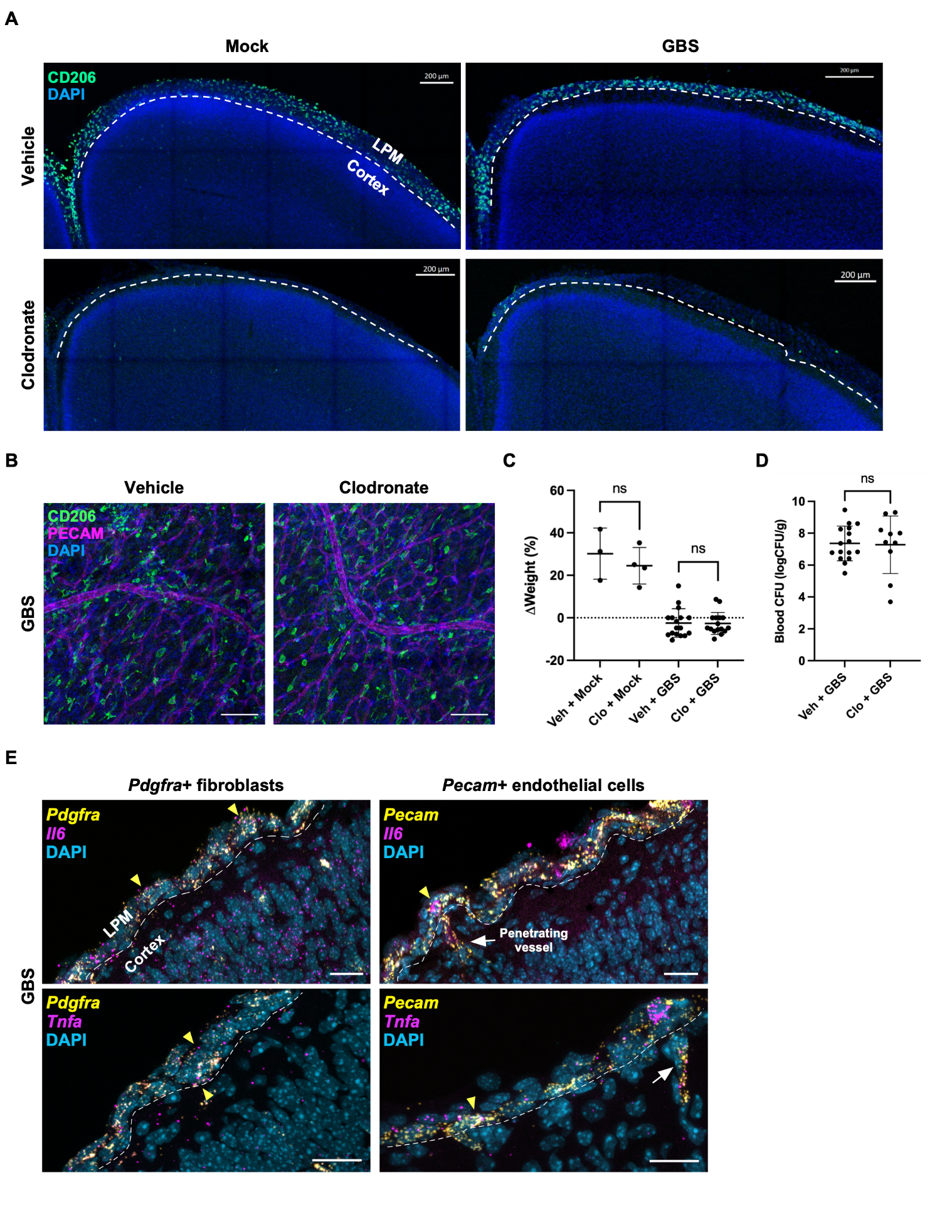

**Supplementary Figure 3. (A)** Visualization of clodronate-mediated L-BAM depletion (24 hpi) in mock- and GBS-infected brains. (100µm coronal brain sections) **(B)** CD206^+^ dural BAMs in vehicle- and clodronate-treated animals. **(C)** % weight change between P2 and P3 and **(D)** blood CFU across vehicle- and clodronate-treated groups. (n=3 vehicle mock, n=4 clodronate mock, n=17 vehicle GBS, n=16 clodronate GBS) **(E)** *Il6* and *Tnfa* transcripts in *Pdgfra*^+^ fibroblasts and *Pecam*^+^ endothelial cells of RNAscope-processed whole head sections (GBS-infected group). Arrowheads indicate *Il6*+ or *Tnfa^+^* cells and arrows indicate penetrating vessels. Statistics: Mann-Whitney *U* test, ns = not significant; mean and SD. Scale bars = (A) 200µm, (B) 100µm, (C) 20µm.

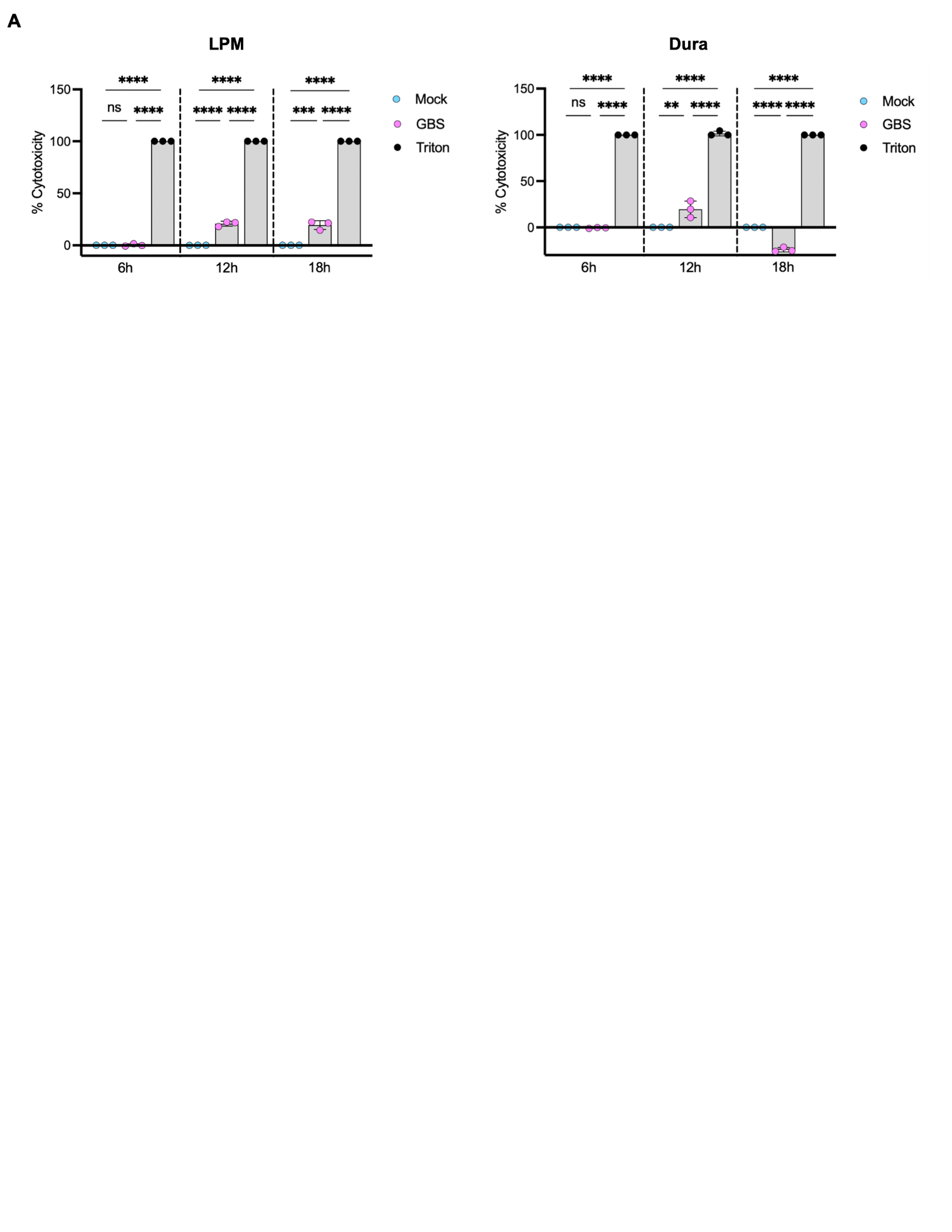

**Supplementary Figure 4.** **(A)** Lactate dehydrogenase (LDH) cytotoxicity assay on primary LPM (left) and dural (right) fibroblasts at 6, 12, 18 hours post-GBS exposure. Maximal LDH activity was quantified by lysing the cells with Triton. (n=3 per group) Statistics: Ordinary one-way ANOVA, * = p < 0.05, ** = p < 0.01, *** = p < 0.001, **** = p < 0.0001, ns = not significant; mean and SD.
